# Gene editing in rat embryonic stem cells to produce *in vitro* models and *in vivo* reporters

**DOI:** 10.1101/112276

**Authors:** Yaoyao Chen, Sonia Spitzer, Sylvia Agathou, Ragnhildur Thora Karadottir, Austin Smith

## Abstract

Rat embryonic stem (ES) cells offer the potential for sophisticated genome engineering in this valuable biomedical model species. However, germline transmission has been rare following conventional homologous recombination and clonal selection. Here we used the CRISPR/Cas9 system to target genomic mutations and insertions. We first evaluated utility for directed mutagenesis and recovered clones with biallelic deletions in *Lef1.* Mutant cells exhibited reduced sensitivity to glycogen synthase kinase 3 inhibition during self-renewal. We then generated a non-disruptive knock-in of *DsRed* at the *Sox10* locus. Two clones produced germline chimaeras. Comparative expression of DsRed and Sox10 validated the fidelity of the reporter. To illustrate utility, oligodendrocyte lineage cells were visualised by live imaging of DsRed in neonatal brain slices and subjected to patch clamp recording. Overall these results show that CRISPR/Cas9 gene editing technology in germline competent rat ES cells is enabling for *in vitro* studies and for generating genetically modified rats.

## Introduction

The rat *Rattus* is a valuable and widely used model organism for studying cognition and behaviour, physiology, toxicology, and various pathologies, such as metabolic and neurodegenerative diseases (Iannaccone and Jacob, 2009). Although the rat was the first mammalian species to be domesticated for biomedical research (Jacob et al., 2010), it has been outpaced in recent years by the mouse, in part because of limitations in directed manipulation of the rat genome. In mice, genome engineering is mostly performed via embryonic stem (ES) cells, and the ease of carrying out such work has been key to their widespread use as an animal model (Capecchi, 2005). Following the definition of culture requirements for mouse ES cells (Ying et al., 2008), rat ES cells have been derived from different rat strains using similar conditions (Buehr et al., 2008; Hirabayashi et al., 2010a; Li et al., 2008). However, rat ES cells are less robust than their mouse counterparts and demand expert handling to maintain robust growth and capacity for germline transmission (Blair et al., 2011), especially after clonal selection required for gene targeting (Hirabayashi et al., 2014; Hirabayashi et al., 2010b; Hirabayashi et al., 2013; Meek et al., 2010; Men et al., 2012; Men and Bryda, 2013; Tong et al., 2010). These technical difficulties have hindered the widespread adoption of rat ES cell transgenesis.

Meanwhile, the development of the CRISPR/Cas9 system (Cho et al., 2013; Cong et al., 2013; Hwang et al., 2013; Ma et al., 2014; Mali et al., 2013; Shen et al., 2013; Wang et al., 2013; Yang et al., 2013) has enabled rat genome editing via direct injection of one-cell embryos (Kim and Kim, 2014; Li et al., 2013a; Li et al., 2013b; Ma et al., 2014; Shao et al., 2014). The injected endonuclease is targeted to a specific DNA sequence by guide RNAs (gRNAs) and introduces double-strand breaks which can be repaired by non-homologous end joining (NHEJ) (Garneau et al., 2010; Lieber, 2010; Marraffini and Sontheimer, 2010). Error-prone NHEJ generally introduces small indels at the cleavage site to generate mutation in one or both alleles of the target sequence. Several knockout (KO) rats have been generated using this method (Li et al., 2013a; Li et al., 2013b). More recently, insertion of large DNA fragments at target loci has been achieved using single-stranded oligodeoxynucleotides (ssODNs) together with CRISPR/Cas9 (Chen et al., 2011; Storici et al., 2006) (Yoshimi et al., 2014; Yoshimi et al., 2016). However, targeting efficiency varies unpredictably between different loci and according to size of the insert. Moreover, both methods are inefficient and require injections of large numbers of embryos with associated maintenance of substantial numbers of animals.

Furthermore, first generation animals are generally mosaic, necessitating additional breeding and genotyping. Therefore this approach does not provide the most efficient use of animals consistent with the 3R principles of reduction, refinement and replacement. CRISPR/Cas9-mediated gene editing has also been applied in spermatogonial stem cells to create knockout rats (Chapman et al., 2015). Germline genome editing can avoid the production of mosaic mutant progeny (Brinster and Avarbock, 1994). However, homologous recombination has yet to be demonstrated, which limits applications.

Here, we tested whether CRISPR/Cas9 technology can be applied efficiently in rat ES cells both for *in vitro* studies and for generation of rats with targeted genomic insertions.

## Results

### Rat embryonic stem cell derivation and culture

The culture conditions for rat ES cells were previously adjusted to reduce spontaneous differentiation by lowering the concentration of the glycogen synthase kinase-3 (GSK3) inhibitor CHIR99012 (CH) (Chen et al., 2013; Meek et al., 2013). However, even under these culture conditions, termed t2iL (see Methods), rat ES cells still exhibit unreliable attachment to feeders, inconsistent growth rate and viability during routine passaging, and a tendency to become tetraploid. These issues pose particular concern during the stringent clonal selection and expansion required for gene targeting. Therefore we assessed several parameters during derivation of new ES cell lines from Dark Agouti (DA) rats in t2iL. Conditions tested were: addition of the PKC inhibitor Gö6983 (Rajendran et al., 2013); addition of vitamin C (250μM) (Esteban et al., 2010); use of Rho kinase inhibitor Y-27632 (Watanabe et al., 2007); substitution of DMEM/F12 with lipid-rich advanced DMEM/F12; reduced oxygen atmosphere. We found that establishment of cell lines was most reliable using advanced DMEM/F12 in the base N2B27 formulation (Ying et al., 2003) together with t2iL, and with addition of Y-26632 in 5% O_2_. We selected one of the newly derived female ES cell lines, DAC27, for use in subsequent experiments.

We first re-tested the effect of the empirical culture modifications on colony formation from single DAC27 cells. Advanced DMEM/F12 and reduced oxygen gave modest but additive improvements (Figure S1). Addition of Rho-kinase inhibitor Y-27632 (Watanabe et al., 2007) had a more substantial effect. The combination of aDMEM/F12, 5% O2 and Y-27632 gave a colony forming efficiency of 82% and moreover made routine passaging more consistent. We therefore incorporated all three modifications into the culture system for targeted genome modification.

### Targeted mutation of *Lef1*

Expression of the canonical Wnt signalling effector *Lef1* has been proposed to underlie the hypersensitivity of rat ES cells to the GSK3 inhibitor CHIR99021 (CH) (Chen et al, 2013). We therefore chose the *Lef1* gene to test the applicability of CRISPR/Cas9 for targeted gene mutation in rat ES cells. A guide RNA was designed using the CRISPR Design tool (http://crispr.mit.edu/) to target the second exon of *Lef1* (Figure 1A).

**Figure 1:**
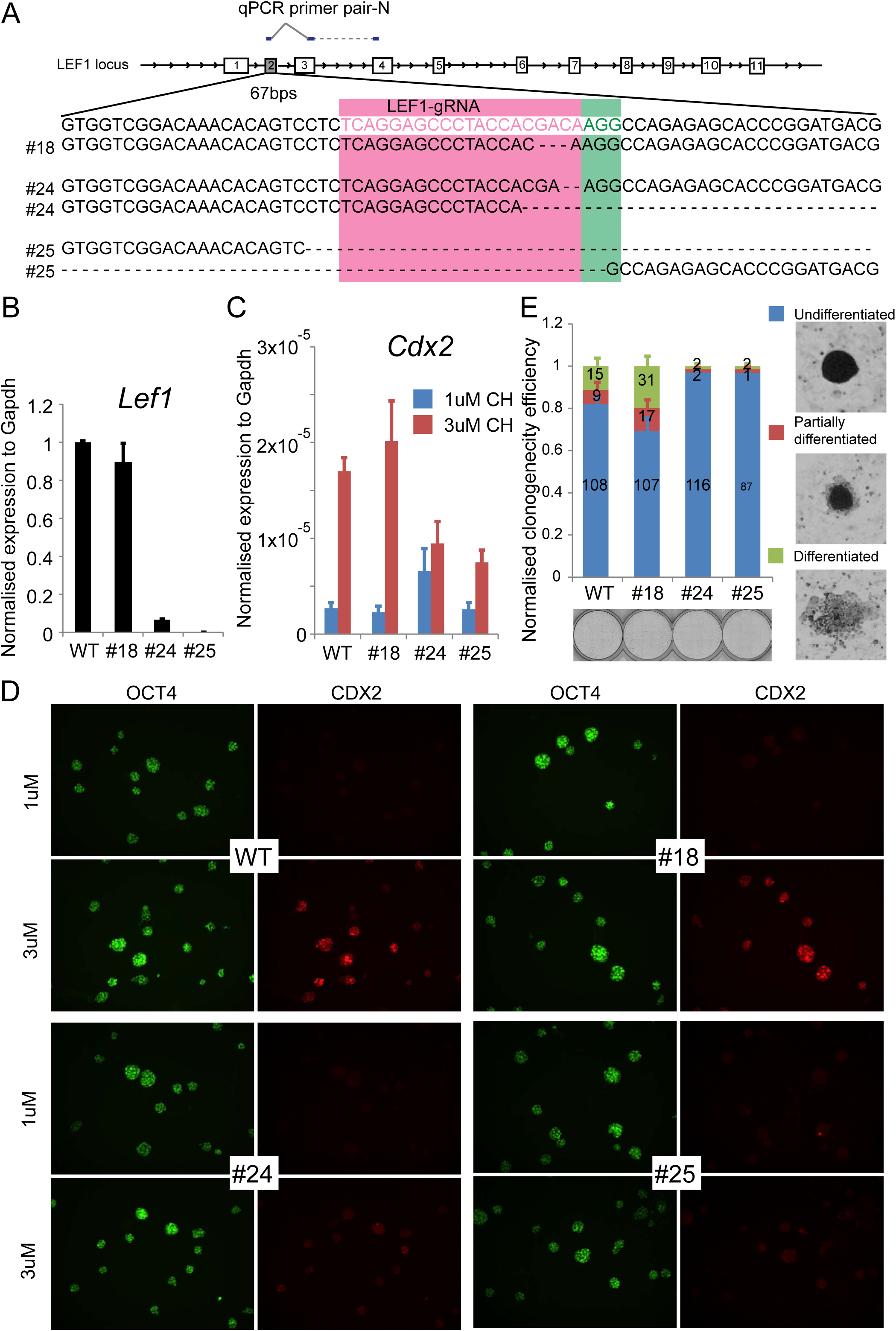
Generation and characterisation of *Lef1* knockout rES cells. (A) Design for CRISPR/Cas9-targeted mutation of *Lef1* exon2. (B) Expression of *Lef1* transcript in parental and *Lef1* mutant rat ES cells assayed by RT-qPCR. (C) Expression level of *Cdx2* transcript in response to CH at 1μM or 3 μM. Error bars represent the standard deviation (STD) of 3 technical replicates. Expression was normalised to *Gapdh.* (D) Fluorescent immunostaining of CDX2 and OCT4. Scale bars represent 100μm. (E) Colony forming assay in the presence of 3μM CH. Error bars represent STD of 3 technical replicates. Undifferentiated, partially differentiated and differentiated colonies were calculated relative to the total number of colonies counted for each line. Scale bars represent 100μm.

For *Lef1* targeting, 1 × 10^6^ rat ES cells were transfected using lipofectamine 2000 with 1.2 μg of expression plasmid containing gRNA and Cas9-2A-GFP. Eight hours post-transfection, cells were replated onto new feeders in fresh medium. Twenty-four hours after replating, GFP-positive cells were sorted by flow cytometry into 10cm culture dishes at a density of 10,000 cells per dish. Fifteen millilitres of medium was added into each dish and no medium change was required thereafter. Five days later, individual colonies were picked, plated into duplicate 96-wells, and expanded briefly before genotyping one of the duplicates.

Genomic PCR followed by gel electrophoresis indicated that five out of thirty-eight (13%) expanded cultures had an overt deletion in one or both alleles of *Lef1* (Figure S2A). We selected three clones with distinct gPCR products: #18, no size change; #24, one smaller band; #25, no wild type band. We subcloned #24 and #25 and repeated the gPCR screen to eliminate the possibility of mixed colonies from the primary plating. We then sequenced the genomic region spanning *Lef1* exon2. Clone #18 had an in-frame deletion of 3 base pairs, with no wild type sequence. Clone #24 had a 2 base pair frame shift mutation in one allele and a deletion of 124 base pairs in the other allele. Clone #25 had deletions of 173 base pairs and 506 base pairs (Figure 1A).

To assess whether *Lef1* expression was indeed disrupted in these three clones we designed primers flanking the gRNA recognition site (Figure 1B). Consistent with an in-frame mutation, clone #18 yielded a PCR product in similar amount to parental cells. Also consistent with sequencing results, clone #25 yielded no detectable product. Analysis of clone #24 on the other hand, indicated a residual level of transcript. This could be due to incomplete nonsense mediated mRNA decay of the frame shifted transcript. We examined LEF1 protein expression by immunocytochemistry using a monoclonal antibody that detects an epitope downstream of the deleted region encoded by exon 2. Strong staining was observed in parental and clone #18 cells, while clone #24 and clone #25 cells were unstained (Figure S2B). Collectively these data indicate that clones #24 and #25 are null mutants lacking LEF1.

We examined the phenotypic consequence of loss of LEF1. In standard 2iLIF medium containing 3μM CH (Ying et al, 2008), expression of Wnt targets related to differentiation, such as CdX2, is appreciable in rat ES cells (Chen et al., 2013; Meek et al., 2013). To investigate whether inactivating Lef1 could alleviate the hypersensitivity of rat ES cells to GSK3 inhibition, we first measured the induction of Cdx2 by RT-qPCR. In parental cells, the expression of Cdx2 increased more than 6 fold when CH concentration was raised from 1μM to 3μM. Clone #18 cells showed a similar response to CH, suggesting that loss of a single amino acid has minor effect on LEF1 function. In contrast, the Cdx2 response to CH was reduced in clones #24 and #25 (Figure 1C). Expression of CDX2 protein was also markedly attenuated in these two *Lef1* mutant clones, as shown by immunofluorescence staining (Figure 1D).

To assess whether loss of Lef1 had an impact on the self-renewal of rat ES cells, we performed colony forming assays in the presence of 3μM CH. Colonies were stained for alkaline phosphatase and scored for level of differentiation, categorised as undifferentiated, partially differentiated, or differentiated. A representative image of each category is shown in Figure 1E. As previously reported (Chen et al., 2013), differentiation was overt in around 20% of parental rat ES cells cultured in 3μM CH. This was also apparent in clone #18. In contrast, in clone #24 and clone #25 mutants fewer than 5% of colonies contained differentiated cells (Figure 1E). We also observed that clones #24 and #25 could be propagated readily in standard 2iL with no evident detriment compared to t2iL, in contrast to parental or clone #18 cells. These results are consistent with LEF1 mediating hypersensitivity of rat ES cells to GSK3β.

### Generation of a non-disruptive *Sox10* knock-in reporter

Based on the proof of principle of genome editing in rat ES cells, we sought to generate a targeted knock-in modification via CRISPR/Cas9 facilitated homologous recombination. We chose the *Sox10* gene in order to create a reporter rat of value to the developmental biology and neuroscience communities. Sox10 is a member of the Sry-related HMG box (Sox) family of transcription factors. It is expressed throughout the developing neural crest (Kelsh, 2006) and in all oligodendroglial lineage cells (Stolt et al., 2002). One particularly attraction of a Sox10 reporter is as a tool for visualising and isolating oligodendrocytes from post-natal animals. Indeed several transgenic mouse lines have been created using the *Sox10* promoter (Kessaris et al., 2006; Rinholm et al., 2011; Shibata et al., 2010; Simon et al., 2012). However, rats in which oligodendrocyte lineage cells are specifically labelled would be a valuable resource due to the relative ease of surgical procedures (Iannaccone and Jacob, 2009) and superiority of demyelinating lesions models in the rat (Woodruff and Franklin, 1999), combined with their greater suitability for learning and cognition assays (Iannaccone and Jacob, 2009).

In common with several other Sox gene family members, Sox10 is haploinsufficient (Kuhlbrodt et al., 1998; Pingault et al., 1998). It is therefore essential to avoid disruption of endogenous Sox10 expression in any knock-in reporter. We designed a construct to insert an internal ribosome entry site (IRES) coupled to a red fluorescent protein (HisDsRed) coding sequence into the 3’ UTR, leaving the *Sox10* gene structure intact (Figure 2A). DsRed is fused to a histidinol resistance enzyme, allowing the potential option for drug selection of Sox10-expressing cells if required. The insertion site was selected five base pairs downstream of the stop codon. Fragments of approximately 1.2 kb of genomic sequence were amplified by genomic PCR to generate 5’ and 3’ homology arms. Sox10 is not expressed in ES cells, therefore positive selection was provided by a PGK-Neo cassette flanked by *Loxp* sites.

**Figure 2:**
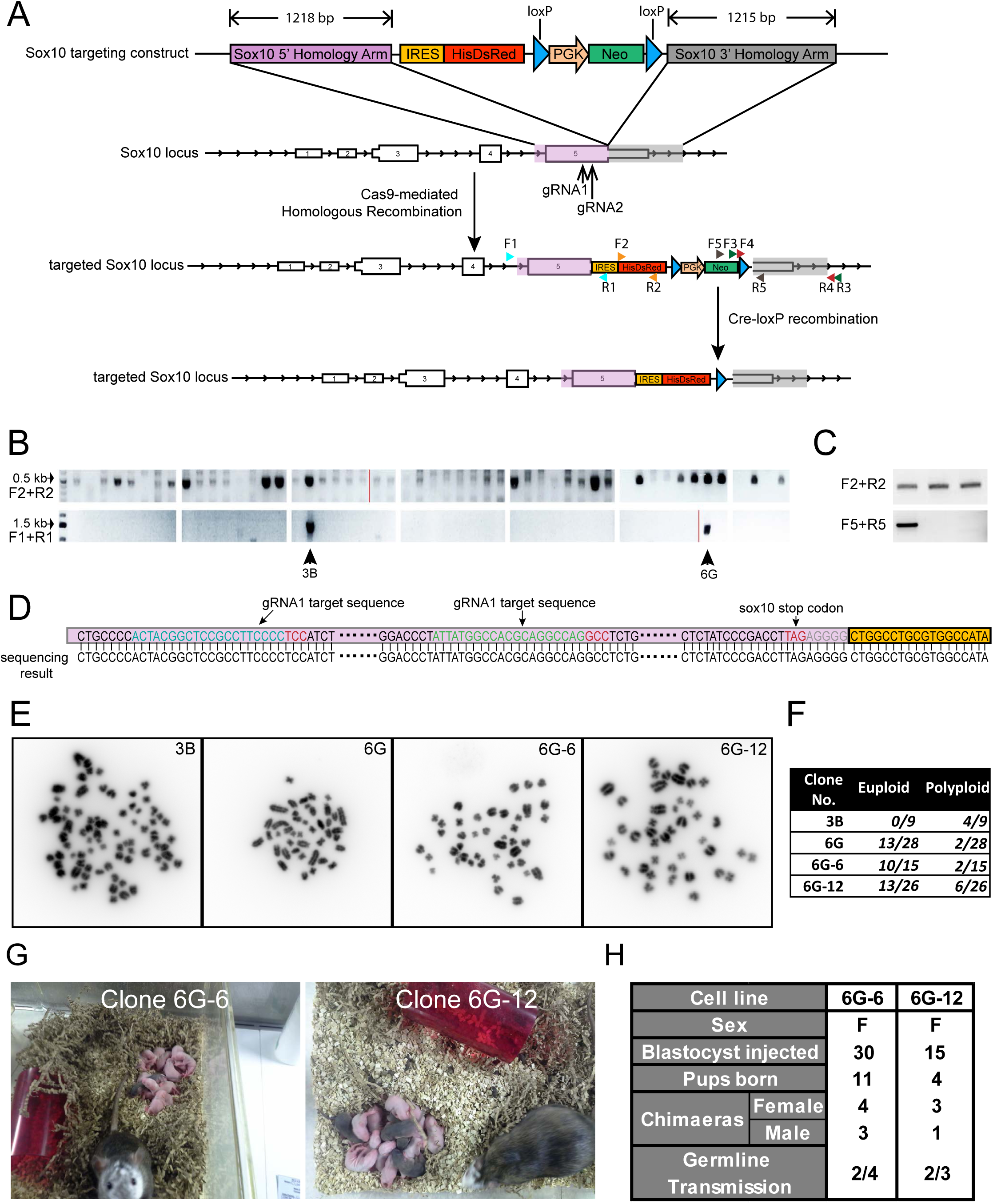
Generation of Sox10-DsRed reporter transgenic rat. (A) Design of *Sox10* targeting. (B) Screen for targeting by genomic PCR. (C) Confirmation of removal of PGK-neomycin selection cassette (D) Genomic sequence around gRNA recognition sites in clone 6G cells. Representative images (E) and chromosome counts (F) of metaphase spreads in Sox10 targeted clones. (G) Chimaeras and germline F1 pups following injection of DA (agouti) ES cells 6G-6 and 6G-12 into SD (albino) blastocysts. (H) Summary of chimaeras and test-breeding.

We designed two gRNAs with recognition sites close to designated insertion site in the 3' UTR. We introduced the gRNAs together with the Sox10-IRES-HisDsRed targeting vector and Cas9 nickase into 1x10^6^ DAC27 rat ES cells via lipofection. Use of Cas9 nickase is reported to increase the ratio of homology-directed repair (HDR) to NHEJ and reduce off-target genome disruption (Cong et al., 2013; Mali et al., 2013; Rong et al., 2014). Transfected cells were replated after 8 hours into 4 x 10 cm dishes on feeders overlaid with Matrigel. After 24 hours G418 (300 μg/ml) selection was applied. Colonies were picked after 7 days of selection and expanded without G418 selection in duplicate for genotyping.

Two out of 52 picked colonies, 3B and 6G, yielded a band of the expected size (∼1.4kb) for homologous recombination detected by genomic PCR using a primer pair flanking the 5’ homology arm (Figure 2B). These two clones were validated further using primers to amplify the reporter region and the 3’ homology arm (Figure S3). To check whether CRISPR/Cas9 editing had created undesired mutations close to the guide RNA recognition sites, we sequenced these genomic regions. No mutations were detected in either clone (Figure 2D).

We assessed the chromosome complement of the two clones by metaphase analysis. Clone 3B contained exclusively hyperdiploid and tetraploid cells (Figure 2D) and was discarded. Clone 6G, had a proportion of tetraploid cells, but 13 out of 28 spreads examined (46%) had a euploid count of 42 chromosomes (Figure 2E). This clone was therefore chosen to proceed to the next step. Cells were transfected with a Cre recombinase expression plasmid and subsequently plated at low density (10,000 cells per 10cm dish) for sub-cloning. Individual colonies were picked and split into duplicate wells of a 96-well plate and cultured with or without G418. Loss of resistance to the antibiotic indicated removal of PGK-Neo cassette, which was confirmed by genotyping (Figure 2C).

Chromosome counts were again checked by metaphase analysis. Two out of twelve clones, clone 6G-6 and clone 6G-12, contained at least 50% euploid cells (Figure 2E-F). These two clones were expanded briefly before injection into blastocysts of the albino SD strain. Coat colour chimaeras were obtained in both cases (Figure 2G). Female chimaeras were test mated to SD males and from each clone two animals proved to be germline competent in the first litter (Figure 2G-H). These data demonstrate that rat ES cells maintained using aN2B27-t2iLY in 5%O_2_ can maintain full competence after two rounds of genetic engineering and clonal selection.

### Sox10 reporter characterisation

Germline offspring from both knock-in clones were bred with SD animals to establish transgenic lines. Heterozygous outcross matings were employed to characterise reporter expression. We evaluated the pattern of DsRed fluorescence at two developmental stages. At embryonic day 11.5 (E11.5), DsRed signal was readily detected in neural crest cell derivatives and the otic placode (Figure 3A), consistent with the pattern of Sox10 expression during embryonic development (Breuskin et al., 2009; Breuskin et al., 2010). At E13.5, fluorescence signals were prominent in dorsal root ganglia (DRG) and trigeminal ganglia, as expected.

**Figure 3:**
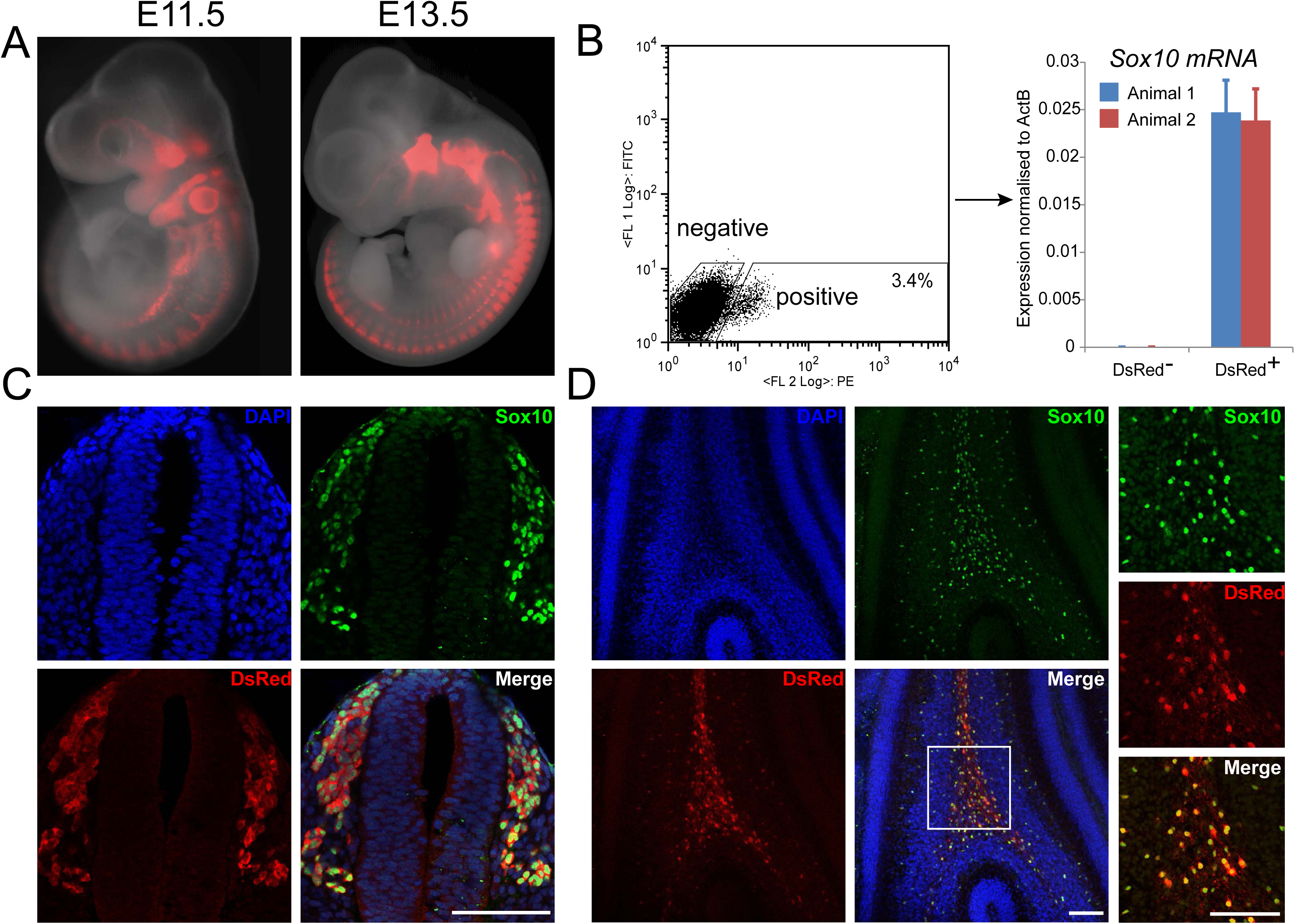
Characterisation of Sox10-DsRed reporter transgenic rat. (A) Fluorescent images of E11.5 and E13.5 embryos. (B) Live sorting of DsRed" and DsRed+ cells from E11.5 embryos followed by RT-qPCR for *Sox10* transcript. Error bars represent STD of 3 technical replicates. (C) Immunostaining for DsRed and SOX10 on 100μm cross-sections of the spinal cord region at E13.5. Scale bars represent 100μm. (D) Immunostaining of 100^,m cross-sections of P7 new born rat cerebellum. The white boxed area is shown in higher magnification in the right panel. Scale bars represent 100μm.

We dissociated E11.5 embryos into single cells and sorted DsRed positive and negative populations by flow cytometry. Approximately 3.5% live cells were positive for DsRed fluorescence. The positive and negative populations were analysed for Sox10 mRNA by RT-qPCR (Figure 3B). Sox10 transcript was only detected in the DsRed positive population, indicating faithful reporting of endogenous Sox10 transcripts. We carried out double immunofluorescent staining for Sox10 and DsRed proteins on sections from E13.5 embryos and P7 new-born rat brain cerebellum. As shown in Figure 3D and E, Sox10 antibody stained nuclei in the DRG region of E13.5 embryos and the cerebellum of P7 neonates. DsRed was detected in the cytoplasm of the same subset of cells. Notably we did not observe expression of DsRed without co-expression of Sox10.

Following outcrossing of F1 animals we set up intercross matings to acquire homozygotes. Homozygous animals were obtained from both clones. We observed no viability or fertility problems in either heterozygotes or homozygotes indicating that Sox10 function is intact. We cryopreserved embryos derived from clone 6G-12 and have deposited live rats with the Rat Resource & Research Center (RRRC).

### Whole cell patch clamp recording from DsRed-positive oligodendrocyte lineage cells

We examined whether this reporter rat can facilitate study of oligodendroglial cells. First we checked whether DsRed signal can be detected in living cells in postnatal rat brain. In freshly prepared coronal brain slices, we identified DsRed^+^ cells by fluorescence microscopy in both cortex (grey matter) and corpus callosum (white matter). Positive cells displayed morphology of oligodendrocyte lineage cells (Figure 4A). All stages of oligodendroglia are expected to be labelled by Sox10::DsRed. To confirm this, fluorescent cells were selected for whole-cell voltage clamp recordings. Cells of different lineage stages were chosen based on morphology assessed through live cell imaging. Once cells were patched in whole-cell mode, lineage stage was further evaluated by morphology through additional Lucifer Yellow dye labelling. The current response evoked by a 10mV pulse (150ms duration) was used to analyse decay constant and input resistance and voltage-current membrane properties used to analyse voltage-gated sodium and potassium currents. Cartoons and live images of cortical oligodendrocyte progenitor cells are shown in Figure 4B. Representative recordings of early oligodendrocyte progenitors, mature oligodendrocyte progenitors and fully differentiated oligodendrocyte are shown in Figure 4C-E. The electrophysiological properties are consistent with the morphological assessments of maturation stage.

**Figure 4:**
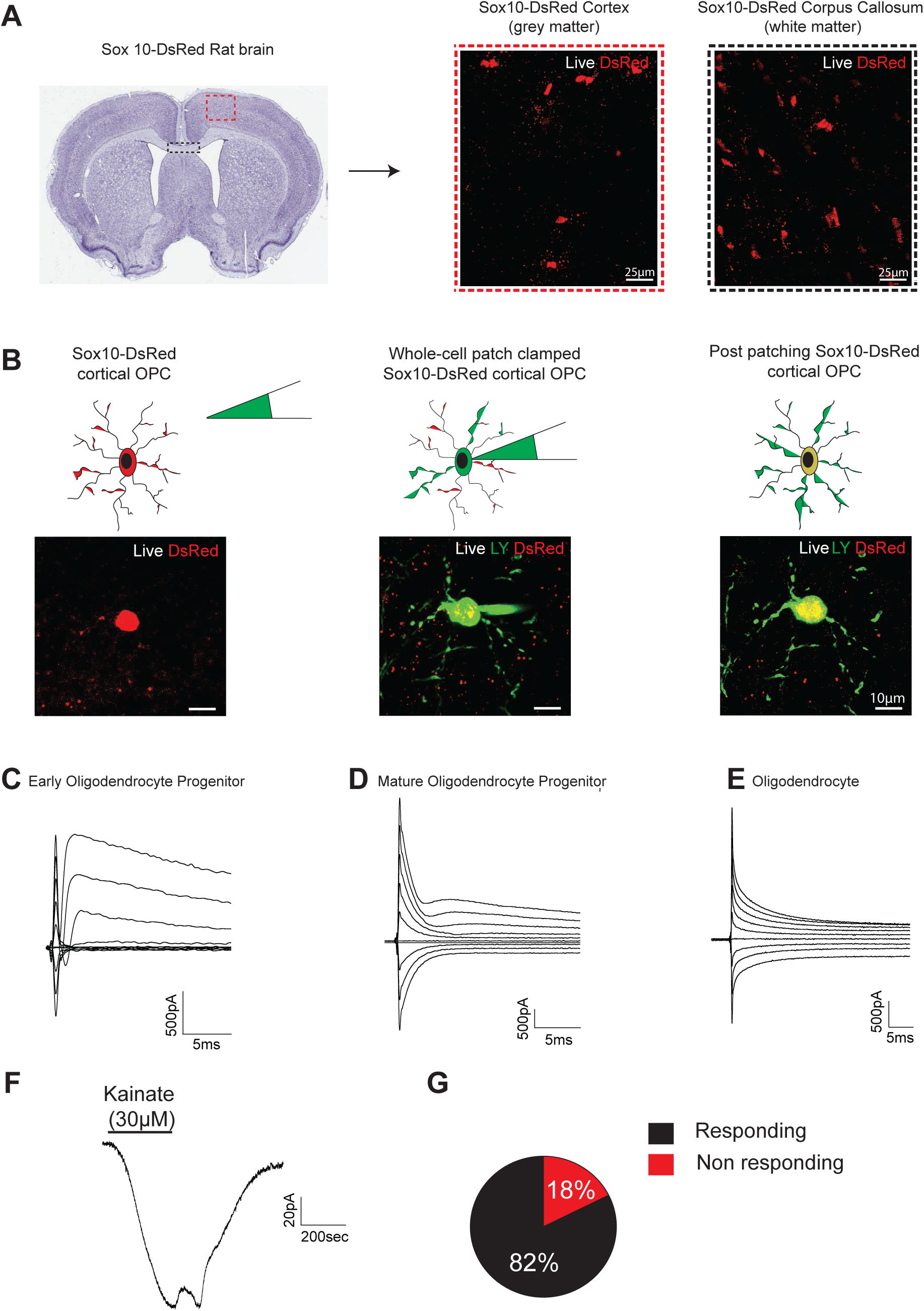
(A) Representation of a cortical Sox10-DsRed rat coronal brain slice. Magnification of red dashed insert shows the live detection of DsRed+ oligodendrocyte lineage cells in the rat cortex. Magnification of black dashed insert represents the live detection of DsRed+ oligodendrocyte lineage cells in the rat white matter (corpus callosum). (B) Schematic and imaging representation of Live DsRed detection with simultaneous dye-filling with Lucifer Yellow (LY) of an oligodendrocyte-lineage cell during whole-cell patch clamp recordings. Characteristic I-V and voltagegated Na_v_ expression levels in DsRed-Sox10+ (C) early oligodendrocyte progenitor cells, (D) mature oligodendrocyte progenitors and (E) oligodendrocytes. (F) Representative response of DsRed-Sox10+ rat oligodendrocyte lineage cells to kainate (30μM) and (G) the percentage of responsive and non-responsive DsRed-Sox10+ rat oligodendrocyte lineage cells to kainate. Total number of cells used for G, n=17 from N=4 separate biological replicates.

Oligodendrocyte lineage cells express glutamate receptors including Kainate receptor (Verkhratsky and Steinhauser, 2000) and respond to Kainate stimulation (Figure 4F). Therefore, we also measured the Kainate response in DsRed+ cells. More than 80% showed a response, further indicating that Sox10::DsRed identifies functional oligodendrocytes in the brain.

## Discussion

Here we have documented the application of CRISPR/Ca9 mediated genome editing in rat ES cells and demonstrated the utility for generation of *in vitro* and *in vivo* models. Incremental refinements of rat ES cell culture conditions conferred more consistent growth and clonogenicity, facilitating recovery of clones after genetic manipulation. In addition, we took two further measures to maximise the probability of germ line transmission: first, we used rat ES cells at low passages; second, we selected diploid clones after each round of clonal selection. Combined with use of the CRISPR/Cas9 system, these refinements increase the practicality of using rat ES cells for gene targeting. Although the incidence of Sox10 knock-in was only 4% of stable transfectants, the construct used lacked any negative selection cassette, commonly included to enrich for homologous recombinants. More importantly, both of the targeted sub-clones selected for blastocyst injection gave robust chimaerism and germline transmission.

The creation of *Lef1* mutant rat ES cells allowed examination of the significance of Lef1 downstream of GSK3 inhibition. This analysis provided further evidence that Lef1 contributes to destabilization of self-renewal via induction of lineage specification genes downstream (Chen et al., 2013; Meek et al., 2013). These results may explain why the optimal concentration of GSK3 inhibitor CH is only 1μM compared with 3μM for mouse ES cells. Interestingly, human naïve pluripotent stem cells also express Lef1 and show a similar requirement for titrated CH (Takashima et al., 2014).

Rat ES cells can be exploited as an *in vitro* differentiation system, complementary to mouse and human pluripotent stem cells. Knock-in reporters are extremely useful tools in this context. For example, the Sox10 reporter generated here could be exploited for monitoring differentiation into neural crest or oligodendroglia, and for purifying desired cell populations.

ES cell-mediated genome engineering has been transformative in mouse genetics and now provides similar opportunities in the rat, which in several areas of physiology and neuroscience has considerable advantages over the mouse as a model species. The *Sox10::DsRed* rat model generated here can facilitate study of neuron-oligodendrocyte interactions and remyelination. Importantly, the rat is preferred to the mouse in this context. Notably rats are used for the cerebellar caudal peduncle (CCP) ethidium bromide model of myelin regeneration (Goudarzvand et al., 2016; Woodruff and Franklin, 1999). The CCP is one of the few fully myelinated tracts in the brain and is often affected by demyelinating disease, such as multiple sclerosis (Preziosa et al., 2014). Rats are used for this lesion because the CCP is not accessible to surgery in mice. More generally, neuropharmacology, cellular distribution of neurotransmitter receptors, and neurotransmitter receptor structure are more similar between humans and rats (Hirst et al., 2003). The Sox10 reporter rat may also be useful in investigations of white matter plasticity, taking advantage of the repertoire of behavioural assays available for rats.

In conclusion, these findings demonstrate that CRISPR/Cas9 methodology can readily be implemented in rat ES cells. Genome editing in rat ES cells constitutes a powerful system for comparative molecular genetic dissection of *in vitro* pluripotent stem cell biology. More broadly, the capacity for germline transmission provides a platform for generating advanced animal models in this important species for biomedical research.

## Experimental procedures

### Cell culture

Rat ES cells were derived from E4.5 blastocysts from the Dark Agouti strain and maintained on γ-irradiated mouse embryo fibroblasts in aN2B27 medium supplemented with t2iL+Y, consisting of MEK inhibitor PD0325901 (1μM), GSK3 inhibitor CHIR99021 (1μM), human recombinant LIF (10ng/mL, prepared in house) and Rho kinase inhibitor, Y-27632 (5μM, Calbiochem). Cultures were maintained in a humidified incubator at 37°C in 7% CO_2_ and 5% O_2_. To prepare aN2B27, we used advanced DMEM/F12 (Gibco™, Catalog no. 12491015). Rat ES cells were routinely passaged by dissociation into single cells with TrypLE™ Express every 48 hours and replated at a split ratio between 1:4 and 1:6. Sufficient culture medium was added when seeding the plates that no change was required until the next passage.

### Gene targeting via CRISPR/Cas9

A newly derived rat ES cell line, DAC27, was used at passage 8 for gene targeting experiments. Guide RNAs were designed using the CRISPR Design tool (http://crispr.mit.edu/) to target desired region (Table S1). For *Lef1* targeting, 1 x 10^6^ rat ES cells maintained in aN2B27 (t2iL+Y, 5% O_2_) were transfected using lipofectamine 2000 with 1.2 μg of expression plasmid containing gRNA, Cas9 and GFP (pSpCas9(BB)-2A-GFP (PX458) was a gift from Feng Zhang (Addgene plasmid # 48138)(Ran et al., 2013). Eight hours post-transfection, cells were replated onto new feeders in fresh medium. Twenty-four hours after replating, GFP-positive cells were FACS sorted onto 10cm culture dishes at a density of 10,000 cells per dish. Fifteen millilitres of medium was added into each dish and no medium change was required thereafter. Five days after seeding, individual colonies were picked, expanded briefly and screened using genomic PCR. For generation of Sox10 knock-in, 1 x 10^6^ cells were transfected with 1.2 μg of gRNA plasmid, 1.2 μg of Cas9 nickase plasmid, and 1.2 μg of targeting vector. Eight hours post-transfection, cells were replated onto 4 x 10 cm culture dishes and G418 selection commenced 24 hours later. Medium was replaced every 24 hours for the first 4 days and every 48 hours thereafter. To ensure robust attachment of rat ES cells during selection a thin layer of matrigel (BD Matrigel™ 1:240 dilution in MEF medium) was applied to the MEF feeder layer 24 hours before rat ES cell seeding. After one week of selection, individual colonies were picked, expanded briefly and screened by genomic PCR. To confirm *Lef1* targeted clones, multiple sets of genotyping primers were used to analyse up to 2.2kb around the gRNA recognition site, and genomic PCR products were sequenced for mutations and deletions (Table S3). To confirm *Sox10* targeted clones, genomic PCR product amplified using *Sox10 set1 (5HA)* primers was inserted into TA clones and sequenced using M13 forward and reverse primers. The region close to *Sox10* gRNA recognition site was also sequenced using customised primer to confirm the absence of mutations.

### Gene expression analysis by quantitative real-time PCR

Total RNA was isolated using the RNeasy Kit (Qiagen) and cDNA prepared using SuperScriptIII (Invitrogen) and 3’RACE adapter primers. Primers and probes used for real-time PCR are listed in Table S2.

### Immunofluorescence staining

Cells were fixed with 4% paraformaldehyde in PBS (pH 7.0) for 30 minutes at room temperature. Subsequently, cells were washed twice with PBST (0.1% Triton X-100 (Sigma) in 1XPBS) and then with blocking solution (4% donkey serum in PBST). Primary antibody solution was prepared by diluting antibody in blocking solution at the concentration listed in Table S4. Cells were incubated with the primary antibody at room temperature for 2 hours or at 4°C overnight, followed by three washes with PBS containing 0.1% Tween 20 prior to incubation with the secondary antibodies at room temperature for 1 hour. After nuclear staining with DAPI (Invitrogen), stained cells were detected by fluorescence microscopy.

### Fluorescence-activated cell sorting

Fluorescent E11.5 embryos were cut into small pieces before incubating in TrypLE™ Express enzyme for 15 minutes at room temperature. Digested tissue was triturated using a p1000 pipette. Enzyme was inactivated and diluted with serum containing wash buffer. Larger debris was removed with 100 μm cell strainers before re-suspension in 1mL of PBS containing 2% BSA for sorting using a Bio-Rad S3™ cell sorter.

### Chromosome analysis

Cells were treated with Colcemide (Gibco, 1:100 dilution) 24 hours after passaging for 2.5 hours. Metaphase chromosome spreads were prepared and imaged at 63x. Chromosomes were counted in discrete spreads.

### Electrophysiology

Parasagittal cerebellar slices (225 μm) were prepared from P3-10 Sox10-DsRed rats using a vibrating blade microtome (Leica VT1200S). After dissection, the brain was placed in a cooled (∼1 °C) oxygenated (95% O2–5% CO2) Krebs solution containing (in mM): 126 NaCl, 24 NaHCO_3_, 1 NaH2PO_4_, 2.5 KCl, 2.5 CaCl_2_, 2 MgCl_2_, 10 D-glucose (pH 7.4). Kynurenic acid was included to block glutamate receptors, which might be activated during the dissection procedure and cause cell damage. During experiments, slices were superfused with HEPES-buffered external solution containing (in mM): 144 NaCl, 2.5 KCl, 10 HEPES, 1 NaH_2_PO_4_, 2.5 CaCl_2_, 10 glucose, 0.1 glycine (to co-activate NMDA receptors), 0.005 strychnine (to block glycine receptors). pH was set to 7.4 with NaOH and solution was permanently bubbled with 100% O2. Recording electrodes were filled with an internal solution comprising (in mM): 130 K-gluconate, 4 NaCl, 0.5 CaCl_2_, 10 HEPES, 10 BAPTA, 4 MgATP, 0.5 Na_2_GTP, 2 K-Lucifer yellow, pH set to 7.3 with KOH; electrode resistance ranged from 5–9 MΩ. Series resistance was left uncompensated and averaged at 30 ± 1.5 MΩ. Electrode junction potential of –14 mV was compensated for. A Multiclamp 700B (Molecular Devices) was used for voltage clamp data acquisition. Data were sampled at 50 kHz and filtered at 10 kHz using pClamp10.3 (Molecular Devices).

## Acknowledgements

We are grateful to William Mansfield and Charles-Étienne Dumeau for generation of chimaeras, Sam Jameson and staff for expert husbandry, Rosalind Drummond for help with qRT-PCR, Yasmin Paterson for help with cell culture, Peter Humphreys for imaging support, Andy Riddell for flow cytometry support, and Marko Hyvonen for recombinant LIF. Y.C. performed and interpreted experiments; S.S. performed the patch clamp recording and S.A. performed live imaging of DsRed+ oligodendrocyte lineage cells and prepared the figure. R.T.K. designed and supervised the electrophysiology experiments. A.S. designed and supervised the study and wrote the paper with Y.C. This research was funded by the European Community project EURATRANS (grant no. HEALTH-F4-2010-241504), by the Biotechnology and Biological Sciences Research Council of the United Kingdom (grant no. BB/H012737/1), the Swiss National Science Foundation programme Sinergia, the Louis Jeantet Foundation, and the Isaac Newton Trust. AS is a Medical Research Council Professor.

